# RID-Rihaczek Phase Synchrony Method Applied to Resting-State EEG: Simultaneous prediction of visuospatial tracking, verbal communication, executive function, neuro-cognitive health, and intelligence

**DOI:** 10.1101/2024.11.07.622537

**Authors:** Toshiya Miyatsu, Arash Mahyari, Jacek Rogala, Timothy Broderick, Peter Pirolli

## Abstract

Brain activity at a resting state and functional connectivity represent individuals’ neural configurations that can be used to predict various performance and trait measures. Yet, most of the previous empirical demonstrations of this phenomena are limited to using resting-state neuroimaging to predict a single task performance/trait measure, and a few studies that predicted multiple task performance/traits measures employed conceptually similar tasks to produce converging evidence for their topic of investigation. In the current paper, we applied RID-Rihazcek phase synchrony, a nonlinear time-frequency-based method for neural computation, to resting-state EEG data and simultaneously predicted a wide variety of measures: standard shooting task performance and novel verbal communication-based team shooting task performance in a simulator, visuo-spatial tracking task performance (Neurotracker), executive function (verbal fluency task), fluid and crystalized Intelligence (Multidimensional Aptitude Battery II: MAB-II), and neurocognitive functioning (Automated Neuropsychological Assessment Metrics-4: ANAM-4) with the average *R*^2^ = .60. Our findings show great promise for RID-Rihazcek phase synchrony and resting-state EEG in general to be used in aptitude assessment.

---

The human brain at rest is not a dormant organ waiting for the next command. Rather, regions of the brain that regularly work together to accomplish tasks show spontaneous coordinated activations even with an absence of an ongoing task, a phenomenon known as functional connectivity^1,2^. Importantly, individuals differ in their brain configurations depending on their genetics^3^ and life experiences^4^, and these individual differences in resting-state connectivity have been successfully used to predict a variety of specific task performance ability^5,6^, traits^7,8^, and even future learning rate^9^.

Among several neuroimaging methods to capture the resting-state brain characteristics, electroencephalography (EEG) emerges as the most accessible and scalable means today. Although there are many demonstrations of resting-state EEG-based performance and trait prediction, the large majority of them are predictions of a single task. A few studies that predicted multiple task performance and traits^5,10^ employed conceptually similar tasks (e.g., visual search task and target shooting) to produce converging evidence for their topic of investigation. Theoretically, however, numerous performance and trait measures could be predicted from the neural configuration characteristics that are extracted from one resting-state neuroimaging data. The omission of this multi-task approach from the literature is especially surprising given that such a method could have significant impact in personnel assessment and selection in a wide variety of areas. In the current study, we employ a high-fidelity computation method for brain connectivity, *RID-Rihaczek Phase Synchrony*, and demonstrate that simultaneous prediction of several task performance, neurocognitive functioning, and trait measures is possible from a single three-minute resting-state EEG reading.

## RID-Rihaczek Phase Synchrony

The methods for computing brain connectivity in EEG are essentially correlation or synchrony measures between different channels of EEG. These connectivity computation methods can be classified into linear or nonlinear approaches^11^. Traditional linear correlation metrics (e.g. Granger causality^12^, directed transfer function^13^, partial directed transfer function^14^) fail to capture the temporal information of EEG signals fully for two reasons. First, these approaches are based on parametric linear auto-regressive models, and therefore susceptible to common problems among parametric models, such as order selection and robustness of parameter estimation under noisy conditions. Second, these models assume stationarity over time and define the transfer function only in the frequency domain and linear interactions between different channels. Such approaches only allow quantification of amplitude effect and fail to separate the effects of amplitude and phase^15^.

Nonlinear phase-synchrony methods address these shortcomings by quantifying time-varying phase synchrony^16–18^. There are two types of commonly used phase synchrony measures. The first type, the Hilbert transform method, bandpath-filters the EEG signal based on target frequency and applies the Hilbert transform to extract the instantaneous phase^19^. However, this method of frequency-dependent phase estimate is indirect and is susceptible to wideband noise. The second type uses the continuous wavelet transform with a complex Morlet wavelet^20^ or the short-time Fourier transform (STFT)^21^ to compute a time-varying complex energy spectrum. While these methods address the stationarity issue, the wavelet transform methods produce biased energy representations and corresponding phase estimates due to nonuniform resolution across time and frequency, and the STFT method has an inherent tradeoff between the resolution of time and frequency due to the window function.

RID-Rihaczek Phase Synchrony, which is based on the Reduced Interference Distribution (RID)-Rihaczek distribution, is a recently developed, enhanced nonlinear time-frequency-based method for neural connectivity computation. It is based on bilinear time-frequency distributions (TFDs) and does not require parameter estimation. RID-Rihaczek Phase Synchrony calculates the synchronization of brain regions in different frequency bands and for all time samples while preventing any aliasing. Preliminary application of this method to EEG data has shown to be less susceptible to volume conduction compared to the traditional continuous wavelet transform estimate of phase difference ^15,22^.

We collected a resting-state EEG reading along with several task performance and trait measures in conjunction with an elite military unit selection process. These measures included a standard shooting task and a novel communication task in a virtual simulator^23^, visuo-spatial tracking task (Neurotracker^24^), executive function (verbal fluency task ^25^), fluid and crystalized Intelligence (Multidimensional Aptitude Battery II: MAB-II ^26^), and neurocognitive functioning (Automated Neuropsychological Assessment Metrics-4: ANAM-4 ^27^). Some of the measures were a part of the standard selection assessment of those undegoing the military selection process (MAB-II and ANAM-4) while the rest were selected to capture important yet often not-assessed traits (communication, visuo-spatial tracking, executive function under stress^28^).

The data processing and predictions were done in the following steps. First, we used the RID-Rihaczek method to compute the phase synchrony for capturing the neural interactions underlying individual differences in task performance and trait measures. For an EEG device with *N* electrodes, the phase synchrony calculations result in *N*(*N*-1)/2 unique connections across *N* different brain regions. Given that different sets of brain connections are related to different performance and trait measures, we selected the connections that had sufficient predictive value for each measure based on the p-value (< .10) of the regression coefficients between the phase synchrony value for each connection and each performance/trait measure. Following previous studies linking similar target measures and resting-state EEG frequency bands^5–7^, we used the phase synchrony values of the beta-2 band (22-29 Hz) for some of the target measures and alpha band (8-12 Hz) for others (see the following section) based on prior literature linking specific bands and trait/performance measures. Lastly, we performed linear multiple regression using the phase synchrony values of the selected connections as the predictors and each performance/trait measure as the dependent variables. Fig. 1 shows the schematics of the data processing and prediction pipeline.

**Fig 1.**
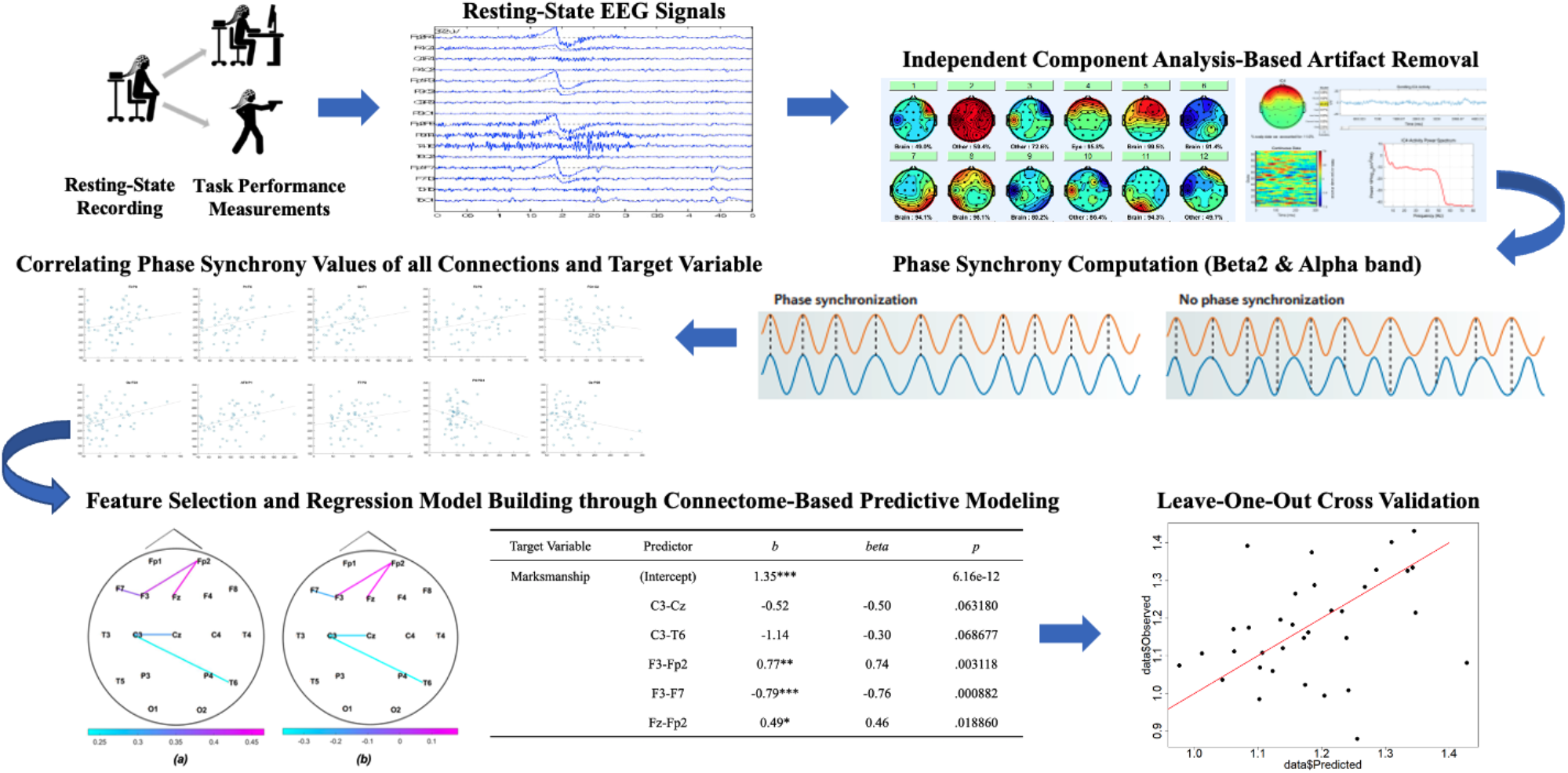
Schematic illustration of the data processing and analysis pipeline.

Using the brain connectivity measures derived from applying the RID-Rihaczek Phase Synchrony as the predictors, we predicted all the significant performance/trait measures mentioned above with a notable accuracy (Average *R*^*2*^ = .62) from a single three-minute resting-state EEG recording. The application of this technique could have wide-reaching implications in personnel selection, assignment, and monitoring.

## Results

We assessed the predictive utility of the features that were derived from applying the RID-Rihaczek Phase Synchrony to the resting-state EEG data in two ways. For each target measure, we present the total variance accounted for and the statistical significance of the final multiple regression model (Table 1). In addition, we present the results of leave-one-out cross validation (LOOCV) in terms of mean square error (MSE) and plot each data point by its predicted value and the actual score (i.e., the ground truth). The list of predictors that were included in the final regression model predicting each target variable, their unstandardized and standardized regression weights, and their *p*-value in the model are reported in Supplemental Material (Table S1).

**Table 1.** Summary measures indicating the fit of the final regression models predicting each target variable.

| Target Variable | $R^2$ | $p$ | LOOCV $MSE$ | $M$ |
| --- | --- | --- | --- | --- |
| Marksmanship | .607 | 4.78e-5*** | 0.02 | 1.17 |
| Teaming Task | .612 | .002936*** | 195.76 | 61.74 |
| Neurotracker | .333 | .01533* | 0.10 | 1.72 |
| Verbal Fluency | .628 | .005594** | 10.35 | 17.90 |
| Performance IQ | .559 | .001239** | 109.71 | 113.06 |
| Verbal IQ | .708 | 3.634e-05*** | 81.28 | 112.22 |
| ANAM-4 | .770 | 1.419e-05*** | 0.56 | 0.01 |

$R^2$ and $p$ are those of the overall model. LOOCV $MSE$ is the mean squared error (MSE) of the leave-one-out cross validation (LOOCV). $M$ is the mean of the target variable. \* $p < .05$ , \*\* $p < .01$ , \*\*\* $p < .001$ .

### Marksmanship task - reaction time (beta-2 band: 22-29 Hz)

Through the feature selection step (see Method for details), connections C3-Cz, C3-T6, F3-Fp2, F3-F7, and Fz-Fp2 are identified as significant. Table 1 shows the results of the connection selection with the values of coefficients for the selected connections and their p-values. The brain topography on the left on Figure 2 a shows these significant connections with their averaged phase synchrony values across all subjects and the middle topography figure shows these connections representing the correlation of the connection with the score.

**Fig 2.**
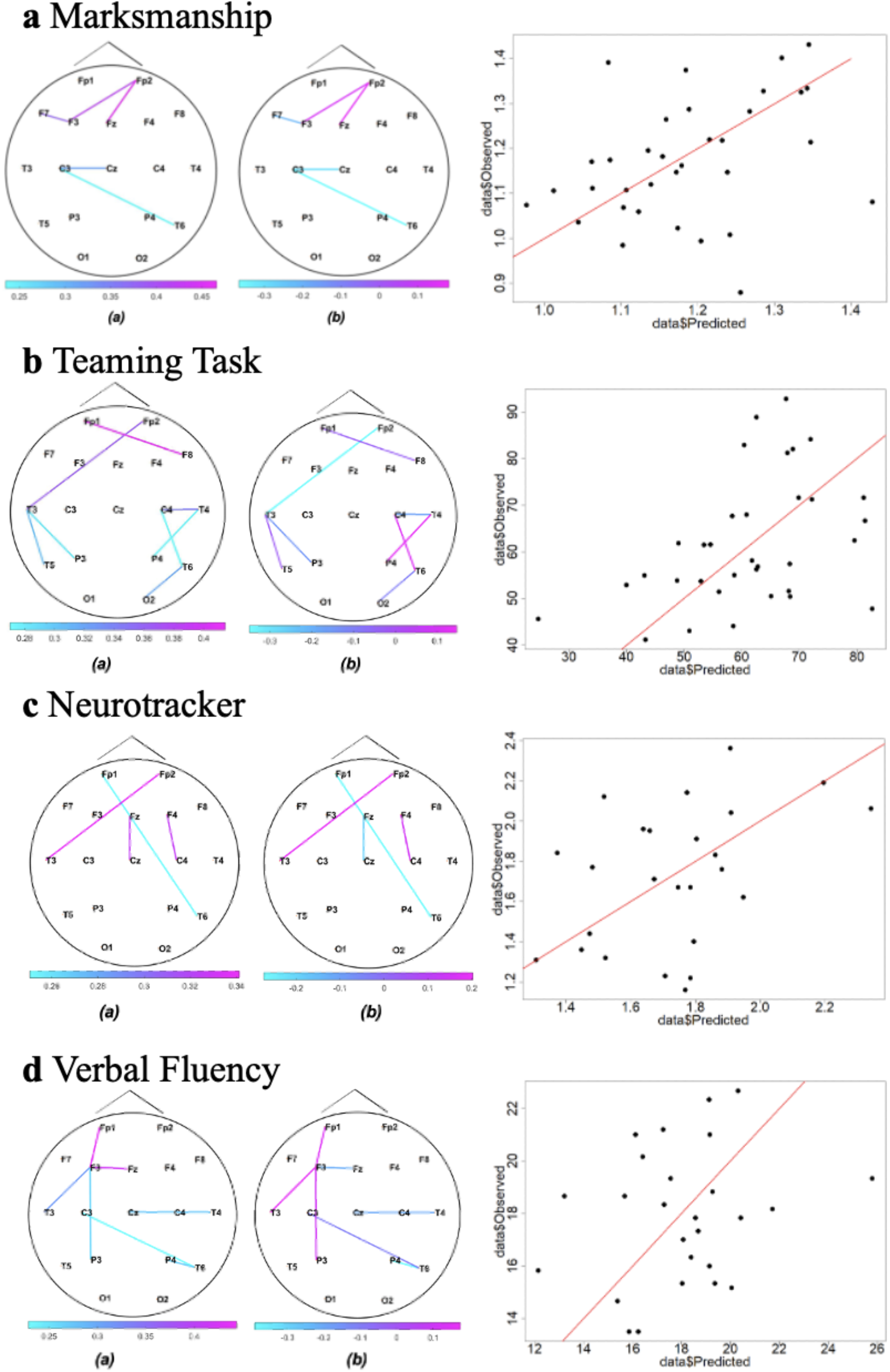
Connectivity maps and LOOCV results for the target variables examined in the beta band. For each panel depicting the results for each target variable (**a**: Marksmanship; **b**: Teaming Task; **c**: Neurotracker; **d**: Verbal Fluency), the topography map on the left (a) shows the average synchrony values across all participants for connections that exceeded the *p* < .10 threshold, the topography map in the middle (b) shows the correlation between the phase synchrony value of the given connection and the target variable, and the scatter plot on the right depicts the results of the LOOCV by plotting the predicted score against the actual score of the given target variable. Each circle represents one subject. The orange line is the optimal prediction line where the predicted score is equal to the observed score.

**Fig 3.**
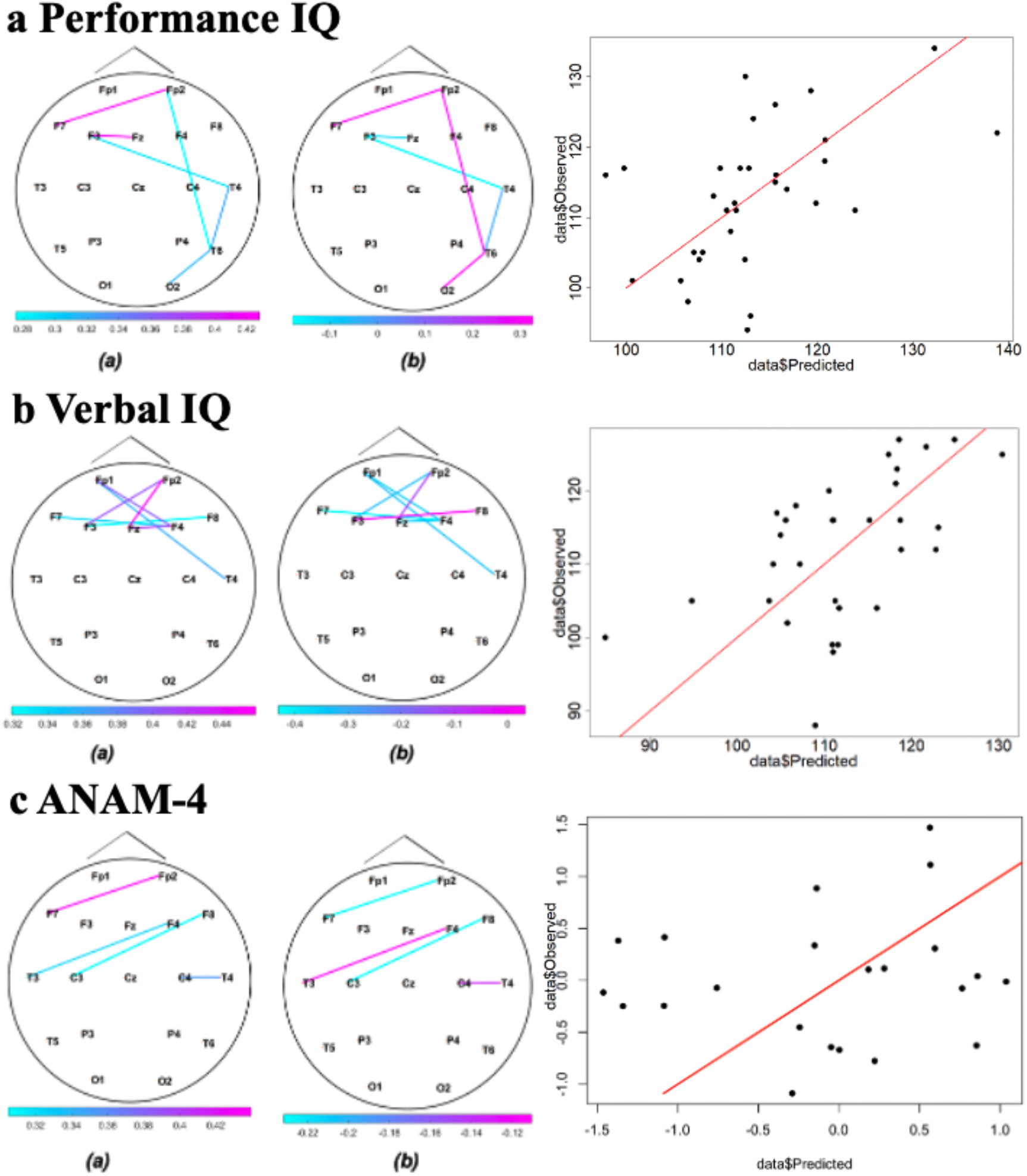
Connectivity maps and LOOCV results for the target variables examined in the beta band. For each panel depicting the results for each target variable (**a**: Performance IQ; **b**: Verbal IQ; and **c**: ANAM-4), the topography map on the left shows the average synchrony values across all participants for connections that exceeded the *p* < .10 threshold, the topography map in the middle shows the correlation between the phase synchrony value of the given connection and the target variable, and the scatter plot on the right depicts the results of the LOOCV by plotting the predicted score against the actual score of the given target variable. Each circle represents one subject. The orange line is the optimal prediction line where the predicted score is equal to the observed score.

We have performed LOOCV procedure for 34 participants, and predicted the average reaction time of the marksmanship task for one holdout participant from a model trained on 33 participants. The MSE of LOOCV for the prediction across all participants is 0.02. The right panel of Figure 2 a shows the scatter plot depicting the predicted scores vs observed scores in the LOOCV.

### Teaming task total completion time (beta-2 band: 22-29 Hz)

Through the feature selection step, connections C3-Cz, C3-T6, F3-Fp2, F3-F7, and Fz-Fp2 are identified as significant. Table 1 shows the results of the connection selection with the values of coefficients for the selected connections and their p-values. The brain topography figure on the left on Figure 2 b shows these significant connections with their averaged phase synchrony values across all subjects, and the middle topography figure shows these connections representing the correlation of the connection with the score.

We have performed LOOCV procedure for 34 participants, and predicted the total completion time for one holdout participant from a model trained on 33 participants. The MSE for the prediction is 195.76. The right panel of Figure 2 b shows the value of the observed scores vs their predicted scores in the LOOCV.

### Neurotracker FinalBL (beta-2 band: 22-29 Hz)

Through the feature selection step, connections Fz-Cz, F4-C4, Fp1-T6, and Fp2-T3 are identified as significant. The brain topography figure on the left on Figure 2 c shows these significant connections with their averaged phase synchrony values across all subjects, and the middle topography figure shows these connections representing the correlation of the connection with the score.

We have performed LOOCV procedure for 25 participants in theNeurotracker FinalBL task, and predicted the total completion time for one holdout participant from a model trained on 25 participants. The MSE for the prediction is 0.10. The right panel of Figure 2 c shows the scatter plot depicting the predicted scores vs observed scores in the LOOCV.

### Verbal Fluency task overall (beta-2 band: 22-29 Hz)

Through the feature selection step, connections P3-F3, C3-T6, F3-Fz, F3-Fp1, F3-T3, P4-T6, and Cz-T4 are identified as significant. Table 1 shows the results of the connection selection. The brain topography figure on the left on Figure 2 d shows these significant connections with their averaged phase synchrony values across all subjects and the middle topography figure shows these connections representing the correlation of the connection with the score.

We have performed LOOCV procedure for 26 participants in the verbal fluency overall task, and predicted the total completion time for one holdout participant from a model trained on 25 participants. The MSE for the prediction is 10.35. The right panel of Figure 2 d shows the scatter plot depicting the predicted scores vs observed scores in the LOOCV.

### Multidimensional Aptitude Battery II (MAB-II) - Performance IQ (alpha band: 8-12 Hz)

Through the feature selection step, connections F3-Fz, F3-T4, Fp2-F7, Fp2-T6, O2-T6, and T6-T4 are identified as significant. Table 1 shows the results of the connection selection. The brain topography figure on the left on Figure 2 e shows these significant connections with their averaged phase synchrony values across all subjects and the middle topography figure shows these connections representing the correlation of the connection with the score.

We have performed LOOCV procedure for 32 participants in the MAB-II performance IQ task, and predicted the total completion time for one holdout participant from a model trained on 31 participants. The MSE for the prediction is 84.65. The right panel of Figure 2 a shows the scatter plot depicting the predicted scores vs observed scores in the LOOCV.

### Multidimensional Aptitude Battery II (MAB-II) - Verbal IQ (alpha band: 8-12 Hz)

Through the feature selection step, connections *F3-Fp2, F3-F8, Fz-F4, Fz-Fp2, F4-Fp1, F4-F7, Fp1-T4* are identified as significant. Table 1 shows the results of the connection selection. The brain topography figure on the left on Figure 2 f shows these significant connections with their averaged phase synchrony values across all subjects and the middle topography figure shows these connections representing the correlation of the connection with the score.

We have performed LOOCV procedure for 32 participants in the MAB-II performance IQ task, and predicted the total completion time for one holdout participant from a model trained on 31 participants. The MSE for the prediction is 81.28. The right panel of Figure 2 f shows the scatter plot depicting the predicted scores vs observed scores in the LOOCV.

### Automated Neuropsychological Assessment Metrics-4 (ANAM-4) (alpha band: 8-12 Hz)

Through the feature selection step, connections *C3-F8, F4-T3, C4-T4, Fp2-F7* are identified as significant. Table 1 shows the results of the connection selection. The brain topography figure on the left on Figure 2 g shows these significant connections with their averaged phase synchrony values across all subjects and the middle topography figure shows these connections representing the correlation of the connection with the score.

We have performed LOOCV procedure for 23 participants in the ANAM-4 task, and predicted the total completion time for one holdout participant from a model trained on 22 participants. The MSE for the prediction is 10.03. The right panel of Figure 2 g shows the scatter plot depicting the predicted scores vs observed scores in the LOOCV.

## Discussion

In a study situated in a selection assessment for a US military unit, we applied RID-Rihaczek phase synchrony to resting-state EEG data and showed that simultaneous predictions of a wide variety of task performance and trait measures were possible from a single 3-minute resting-state EEG recording. We have created a unique resting-state EEG data set accompanied by resting-state ECG ^29^, EOG, and various performance/trait measures. Our findings, both the individual pieces that link certain brain connectivity characteristics with a given performance/trait measure as well as the aggregation of these pieces, have various basic and applied implications.

Previously, the mechanisms through which the linkage between resting-state neuroimaging characteristics and task performance arise were discussed not only in terms of the trait reflected in the functional connectivity, but also in terms of the mental state reflecting the better preparedness of those who are trained in or familiar with the task^6^. Such explanation is plausible from the data obtained in typical resting-state EEG experiments where the participants complete resting-state EEG recording and known tasks in succession in a single experimental session. However, various task performances in our study were collected before and after the resting-state EEG recording without notifying the participants what they would be tested on (see Method section below). Thus, the current findings add to the growing literature indicating that one’s unique neural configurations can be measured through resting-state EEG, and such individual differences underlie various levels of task performance and trait measures.

While the simultaneous predictions of the performance/trait measures include meaningful replications of previous studies, such as the associations between the beta band connectivity and marksmanship^5,6^ as well as the alpha band connectivity and intelligence^7,8^, they also include novel findings. To our knowledge, this is the first study linking EEG connectivity measures with a verbal communication-based teaming task, visuospatial tracking (Neurotracker), verbal fluency task, and neurocognitive health (ANAM-4). Despite the novel application of RID-Rihaczek phase synchrony, some of these findings are consistent with previous findings using other methods of computing brain connectivity. For instance, the novel teaming task involves visual processing of geometric shapes, memorizing complex task order, and listening to the partner and integrating information from multiple sensory sources^30^. Indeed, the channels that were predictive of the teaming task performance included various occipital, temporal, and parietal channels that have been implicated in these types of information processing and task execution^31,32^. Furthermore, the left hemisphere dominance, especially in the temporal and frontal regions, of the connections underlying good verbal fluency performance is consistent with numerous fMRI studies implicating the importance of the left inferior frontal gyrus (LIFG) in this task^33^, and the successful prediction of ANAM-4 performance which is a reaction time-based assessment of neurocognitive health from the alpha band is consistent with previous findings associating the alpha band resting-state activities with attentional capacity^34,35^.

In general, our findings are consistent with the notion that resting-state alpha and beta band activities reflect attention modulation and higher order cognitive processes, such as the cingulo-insular-thalamic network-mediated maintenance of tonic alertness^36^, and motor learning^37^. The current findings, thus, extend the previous findings linking the activities in the alpha and beta bands with attentional investment at rest^38^ and emphasize the importance of these bands in resting-state EEG-based performance/trait measure predictions.

One unique aspect of our procedure is that we collected the resting-state EEG right after the special force unit selection assessment week was over. To our knowledge, this is the first empirical demonstration of the predictive utilities of resting-state neuroimaging data that were collected when subjects were *not* well-rested. The applied nature of the current study enabled the exploration of the relationship between resting-state EEG characteristics and numerous performance/trait measures but also constrained the condition in which we collected the resting-state data. Although the test-retest reliability of resting-state EEG is generally good^39–41^, it has also been shown that exposure to acute^42,43^ stressful events as well as fatigue influence resting-state EEG characteristics significantly. Thus, it is possible that collecting resting-state EEG in this condition somehow affected, or even enhanced, the predictive utility of the resting-state EEG data. Our findings warrant future work aimed at identifying ideal conditions in which resting-state EEG data are obtained to maximize their predictive utilities. In doing so, researchers should consider not only ordinary states, but also conditions known to reliably produce certain psychological and neural states, such as post-stressful events or cold exposure. Such investigation can inform optimal resting-state EEG data acquisition conditions, stability of the predictive utilities of resting-state EEG across distinct psychological states, and performance readiness as well as stress tolerance measurement through resting-state EEG.

The findings discussed above are based on a relatively small sample size (N = 34), and thus, should be taken with caution as a proof of concept of using resting-state EEG and RID-Rihaczek phase synchrony as a multipurpose assessment tool. However, the current study is the first empirical demonstration showing that simultaneous predictions of multiple, distinct performance/trait measures is possible from a brief task-free EEG recording, and thus it should pave the way for developing a new kind of cognitive testing. The strength of this approach lies in its versatility and growth potential. Besides the performance/trait measures and the brain connectivity features we have covered, there are numerous basic research linking resting-state EEG features with neurocognitive and psychopathological conditions, such as Alzheimer’s disease^44–46^, anxiety disorder^47,48^, depression^49,50^ and antidepressant treatment response^51^, as well as other performance and learning measures, including decision making^52^, sustained attention^53^, and motor skill acquisition^54^. As the basic science advances and more validation research is done, one can continuously add algorithm pipelines to extract new predictive features to their system. Importantly, these future additions can be applied to the existing resting-state data without the need for additional assessment, and the predictive accuracy of the algorithms will only increase as more data are acquired. Such technology, if realized, could provide a one-stop assessment of specific task performance, learning capability, general aptitude, and neurocognitive health, and have wide-reaching applications.

## Methods

### Participants

Thirty-four male US military officers (*M*_*age*_ = 25, *SD*_*age*_ = 4.56) participated in the study. They were recruited from among the candidates going through a week-long selection assessment process at a US military unit who had provided consent prior to the selection week and remained on-site until the end of the selection week.

### Materials

#### Marksmanship task

The Marksmanship task is a range shooting task implemented in the virtual shooting simulator VirTra^55^. The participants were presented with 30 targets (15 shoot and 15 no-shoot targets) per round in a virtual shooting range one by one at a 1.5 second interval. Depending on whether the shoot target was hit and the shots’ centrality, each shot received 3, 2, 1, or 0 points. All participants completed 2 rounds of the marksmanship task. Average reaction time and total score were calculated and used as the measures of interest.

#### Virtual Integrated Social Task (VISTa)

VISTa is a verbal communication-based shooting task which was implemented in VirTra and designed to measure communication, teamwork, and fluid reasoning under time pressure. Two participants take the role of a leader and a follower separated by a divider. Both are presented with three images of geometric shapes moving vertically. Only the leader has the target highlighted in red, and he has to describe the target to the follower. Upon receiving the description of the target, the follower shoots the target on his screen, confirms the hit, and the leader shoots an advance target to advance to the next trial. Good communication advances the trials quickly. Crucially, the images were designed in a way that specification of multiple features (e.g., color, size, tilt, etc.) is necessary to identify the target, such that they elicit back-and-forth communication. There were 4 trials in 1 round. Each participant completed 4 rounds: 2 as the leader and 2 as the follower. Several performance metrics from VISTa predicted communication, teamwork, and leadership demonstrated during the selection week^30^. The average completion time was calculated and used as the measure of interest.

#### Neurotracker (NT)

NT is a 3D multiple object-tracking (MOT) task designed to measure visuo-spatial tracking, peripheral vision, and visual information processing. It has previously been shown to predict a variety of sports performance expertise^24^. For each trial, participants were presented with 8 spheres in their 3D visual field, and tracked the movement of 4 of them for 8 seconds. The speed of the spheres’ movement in the next trial was adjusted based on the success/failure to track all 4 target spheres. Each session included 20 trials, and all participants were requested to complete 21 sessions spread across at least 7 days as a part of cognitive testing prior to the arrival to the selection assessment location. Due to logistical limitations (e.g., shipping equipment such as 3D glasses, non compliance, etc.), participants varied in the number of sessions they completed. The final speed at which a given participant could track all 4 target spheres (visual tracking speed: VTS) was used as the measure of interest.

#### Verbal fluency task (VF)

The VF task is an established measure of executive function^56^. It was administered as a part of a speech-based cognitive testing^28^. The average number of words recalled across 6 standard categories (body parts, fruits, animals, words starting with the letter “A”, “F”, and “C”) was used as the measure of interest.

#### Multidimensional Aptitude Battery II (MAB-II)

MAB-II is a battery of 10 tests designed to measure intelligence^26^ and among the most commonly used cognitive tests in the US military. Corresponding to the well-validated concepts of fluid intelligence and crystalized intelligence^57^, it calculates the performance IQ and the verbal IQ, and they were used as the measures of interest.

#### Automated Neuropsychological Assessment Metrics-4 (ANAM-4)

ANAM-4 is a test of neurocognitive functioning designed to detect cognitive abnormality, such as traumatic brain injuries^58^. Participants completed a set of 10 tasks from the core battery that were designed to measure attention, concentration, reaction time, memory, processing speed, and decision-making. Normalized average of the median reaction time from all tasks (overall test battery mean: OTBM^59^) was calculated and used as the measure of interest.

### Experimental procedures

The data were collected in conjunction with a week-long selection assessment process at a US military unit. Fig 4 shows the timeline. The selection week consisted of a series of exercises that challenged the participants physically, mentally, and emotionally. Prior to the arrival at the selection assessment site, participants completed a pre-selection remote assessment which included Neurotracker and the verbal fluency task. MAB-II and ANAM-4 were collected as a part of the selection week assessment. After the selection process was completed, there was an assessment day where the resting-state EEG, VirTra marksmanship, and VISTa data were collected among other assessments (e.g., physical tests such as strength, endurance, and cardiovascular capacities which are not reported in the current paper).

**Fig 4.**
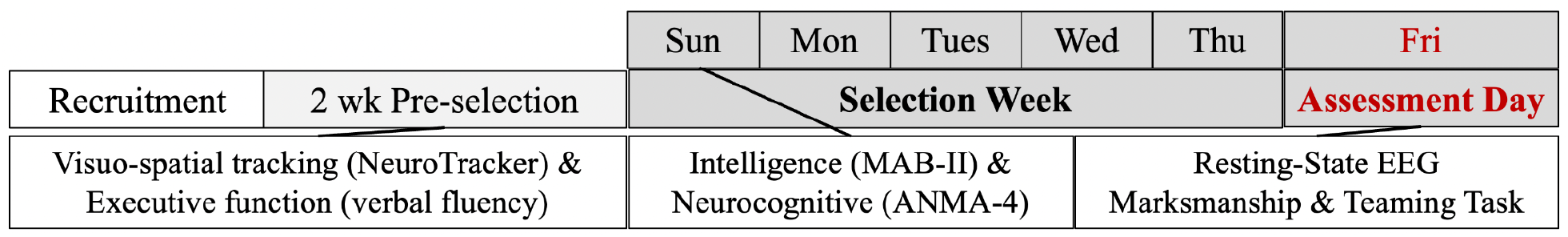
A schematic illustration of the time course of the data collection.

### Electrophysiological data

#### Recording and preprocessing

Three-minutes eyes-closed resting-state EEG were collected at a sampling rate of 300 Hz using a dry electrode 21-channel headset (Wearable Sensing, DSI-24) with the 10-20 system positioning. The Pz electrode was used as the reference electrode and subsequently excluded from the final analysis along with A1 and A2 which served as back-up reference electrodes, leaving 18 channels for the following analysis.

The acquired data were preprocessed using the open source EEGLAB toolbox^60^. The preprocessing pipeline included 1.0-50.0 Hz bandpass filtering, manual exclusion of noisy parts of the data, and removal of eye movement and muscle artifacts using independent component analysis (ICA). Six participants had a channel that had low signal quality, and these subjects’ data were analyzed with 17 channels.

### Phase Synchrony Computation

The connectivity is quantified using a recently introduced phase synchrony measure based on Reduced Interference Distribution (RID)-Rihaczek distribution^15^. The first step in quantifying phase synchrony is to estimate the time and frequency dependent phase, *Φ*_*i*_(*t, ω*), of a signal, *s*_*i*_,is 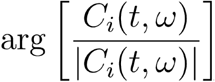, where *C*_*i*_(*t, ω*) is the complex RID-Rihaczek distribution. The details of the RID-Rihaczek distribution and the corresponding synchrony measure are given in our previous publications^15,61,62^.

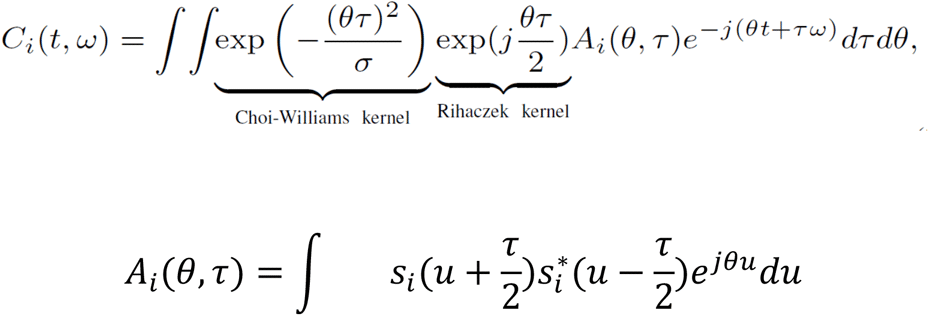

Where *A*_*i*_(*θ, τ*) is the ambiguity function of the signal *s*_*i*_. The phase synchrony between channels *i* and *j* at time *t* and frequency*ω* is computed using Phase Locking Value (PLV):

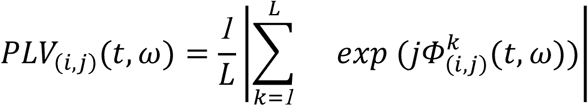

where *L* is the number of trials in the ERP measurements and 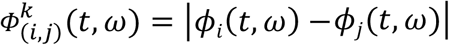 is the phase difference between the two channels for the *k*th trial. The single-trial phase differences, represented by complex numbers, are averaged across trials. The absolute value of this complex average yields the magnitude value that defines the phase locking value. A phase locking value of 0 indicates completely uniform random phase angle differences across trials, whereas phase locking value of 1 indicates completely consistent phase angle differences across trials. This measure quantifies the extent to which oscillation phases are similar across different electrodes at each time-frequency point.

### Regression Model

The phase synchrony calculated for N electrodes results in 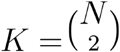 unique values. Not all values are strongly correlated with the task under study. Thus, we perform statistical significant testing to select PLVs highly correlated with the task. Our goal is to predict the score of the task from the PLVs of the resting states EEGs. Our proposed approach is an extension of connectome-based predictive modeling (CPM)^63^. We formulate the task score prediction as *g(y) = αX + β*, where *y* is the participant’s score, *g*(·) is the link function, *X* is the vector of all 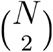 unique PLVs, *α* is a -dimensional unknown vector parameter, and *β* is an unknown constant estimated from the training data. Once the model is established, we draw inference about each estimated parameters in *α* via statistical testing. More specifically, our hypothesis testing is:

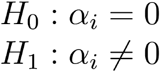

Because the proposed method is a multiple regression model with *K* + 1 unknown parameters, +we perform a forward stepwise inference to test the hypothesis testing for each added parameter *α*_*i*_. Our final results is a multiple regression model with a subset of *α*_*iS*_ that will be used in the final testing and evaluation.

## Data Availability

Data will be made available by request and upon completing required paperwork.

## Acknowledgements

This work was supported by DARPA/AFRL Contract FA8650-19-C-7944 within the DARPA Measuring Biological Aptitude (MBA) program. All statements of fact, opinion or conclusions contained herein are those of the authors and should not be construed as representing the official views or policies of DARPA, AFRL or the U.S. Government. Authors thank Marcas Bamman, 7Art Finch, Alex Oliver, Kurtis Gruters, Ian Perera, Mike McCullough, Kody Coleman, Shelby Greene, Vanessa Oviedo, Andrew Koutnik, and other members of the PEERLESS team for their assistance with data collection. This document was approved by DARPA for Public Release, Distribution Unlimited.

## Supplemental Materials

**Table S1.**
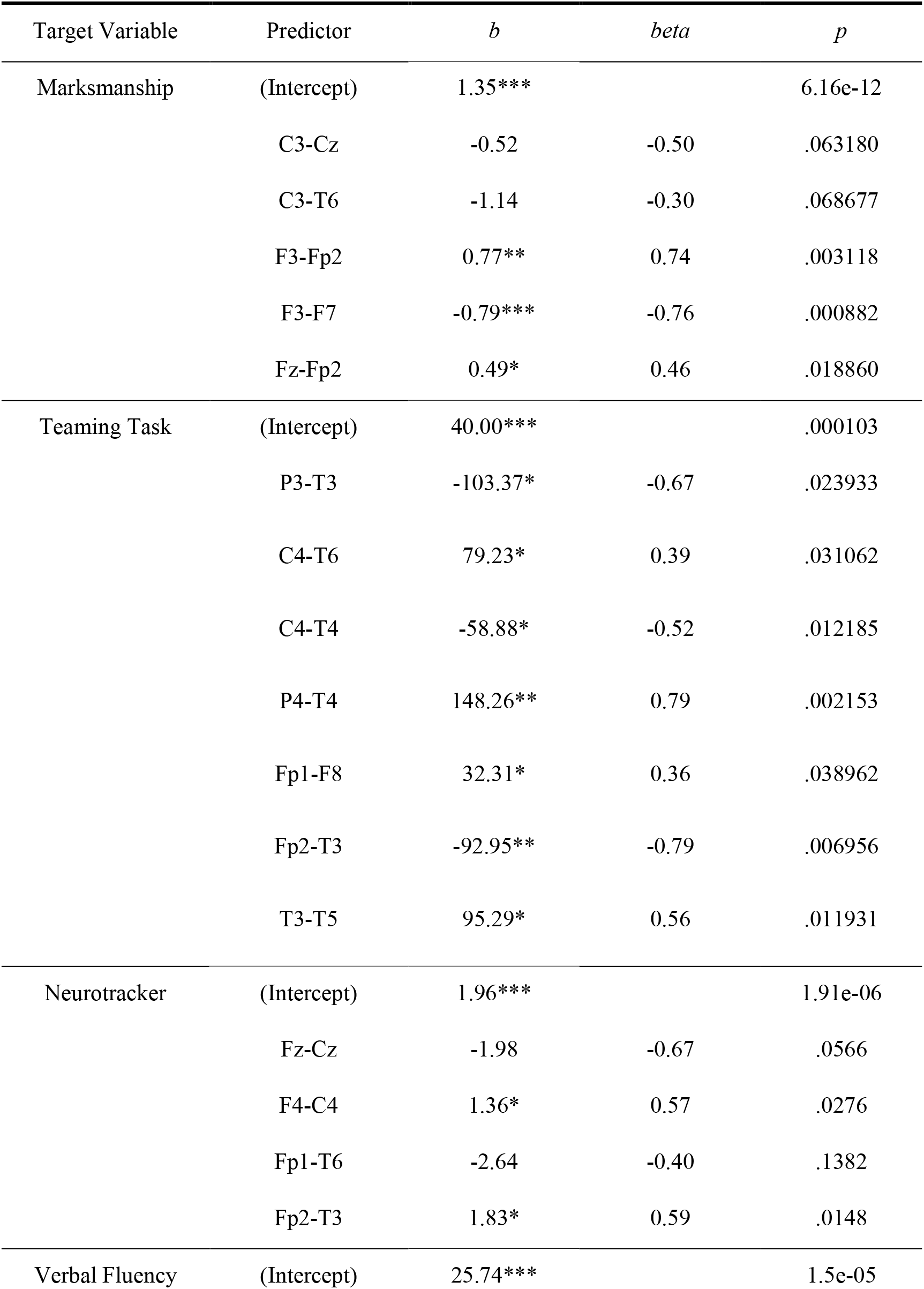
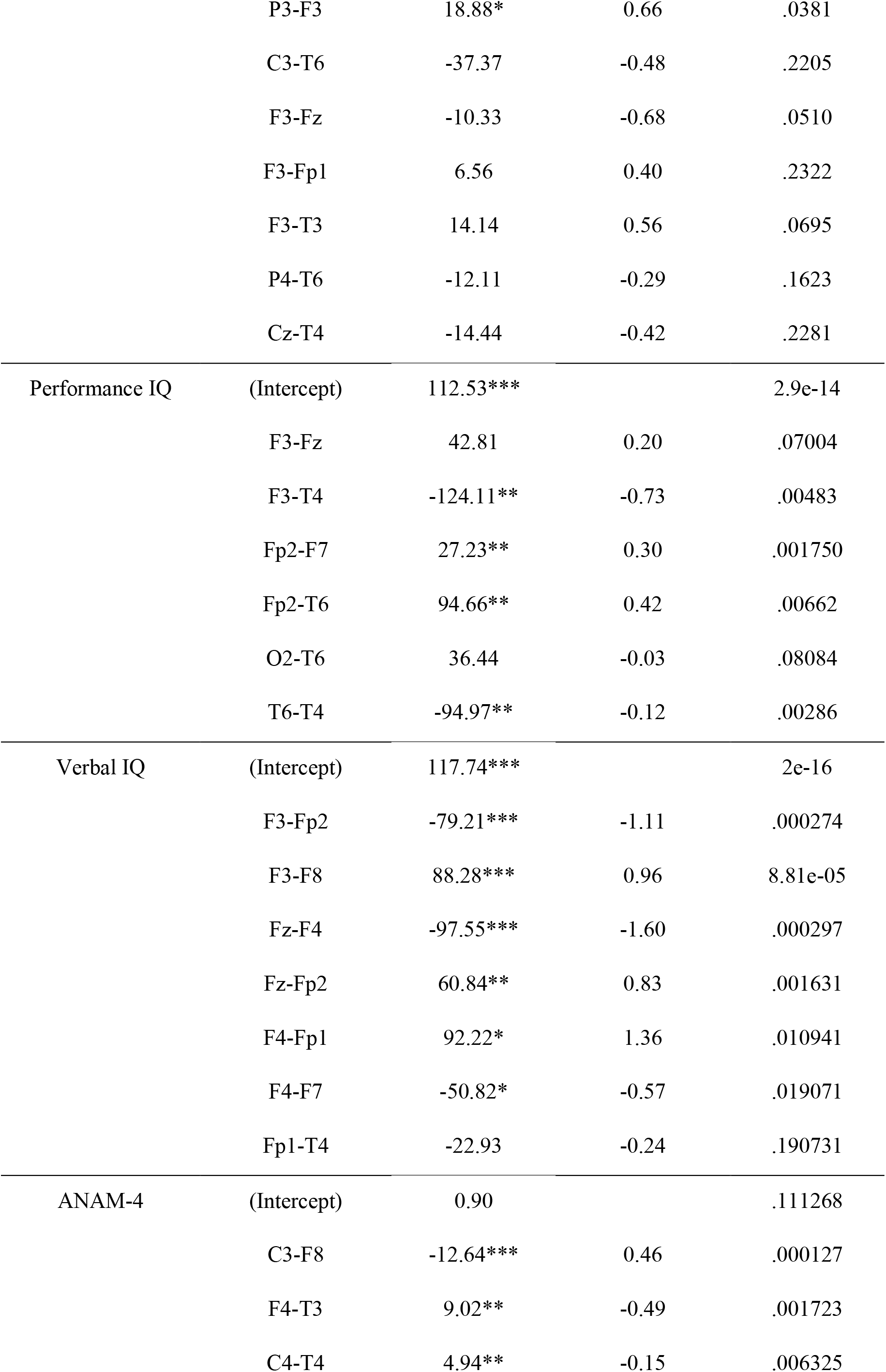
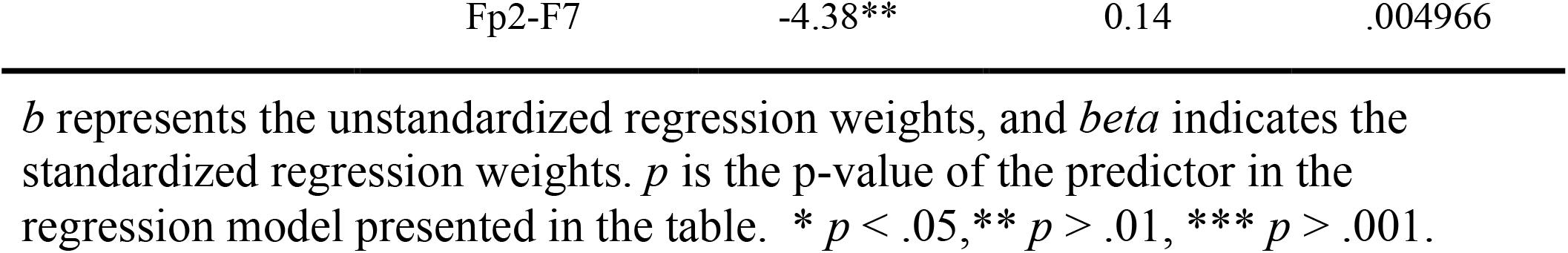
The list of predictors that were included in the final regression model predicting each target variable, their unstandardized and standardized regression weights, and their *p*-value in the model.

