## Supplementary Material for "RID-Rihaczek Phase Synchrony Method Applied to Resting-State EEG: Simultaneous prediction of visuospatial tracking, verbal communication, executive function, neuro-cognitive health, and intelligence"

**Supplemental Materials**

**Table S1.** The list of predictors that were included in the final regression model predicting each target variable, their unstandardized and standardized regression weights, and their *p*-value in the model.

| Target Variable | Predictor | *b* | *beta* | *p* |
|---|---|---|---|---|
| Marksmanship | (Intercept) | 1.35*** | | 6.16e-12 |
| | C3-Cz | -0.52 | -0.50 | .063180 |
| | C3-T6 | -1.14 | -0.30 | .068677 |
| | F3-Fp2 | 0.77** | 0.74 | .003118 |
| | F3-F7 | -0.79*** | -0.76 | .000882 |
| | Fz-Fp2 | 0.49* | 0.46 | .018860 |
| Teaming Task | (Intercept) | 40.00*** | | .000103 |
| | P3-T3 | -103.37* | -0.67 | .023933 |
| | C4-T6 | 79.23* | 0.39 | .031062 |
| | C4-T4 | -58.88* | -0.52 | .012185 |
| | P4-T4 | 148.26** | 0.79 | .002153 |
| | Fp1-F8 | 32.31* | 0.36 | .038962 |
| | Fp2-T3 | -92.95** | -0.79 | .006956 |
| | T3-T5 | 95.29* | 0.56 | .011931 |
| Neurotracker | (Intercept) | 1.96*** | | 1.91e-06 |
| | Fz-Cz | -1.98 | -0.67 | .0566 |
| | F4-C4 | 1.36* | 0.57 | .0276 |
| | Fp1-T6 | -2.64 | -0.40 | .1382 |
| | Fp2-T3 | 1.83* | 0.59 | .0148 |

| | | | | |
|---|---|---|---|---|
| Verbal Fluency | (Intercept) | 25.74*** | | 1.5e-05 |
| | P3-F3 | 18.88* | 0.66 | .0381 |
| | C3-T6 | -37.37 | -0.48 | .2205 |
| | F3-Fz | -10.33 | -0.68 | .0510 |
| | F3-Fp1 | 6.56 | 0.40 | .2322 |
| | F3-T3 | 14.14 | 0.56 | .0695 |
| | P4-T6 | -12.11 | -0.29 | .1623 |
| | Cz-T4 | -14.44 | -0.42 | .2281 |
| Performance IQ | (Intercept) | 112.53*** | | 2.9e-14 |
| | F3-Fz | 42.81 | 0.20 | .07004 |
| | F3-T4 | -124.11** | -0.73 | .00483 |
| | Fp2-F7 | 27.23** | 0.30 | .001750 |
| | Fp2-T6 | 94.66** | 0.42 | .00662 |
| | O2-T6 | 36.44 | -0.03 | .08084 |
| | T6-T4 | -94.97** | -0.12 | .00286 |
| Verbal IQ | (Intercept) | 117.74*** | | 2e-16 |
| | F3-Fp2 | -79.21*** | -1.11 | .000274 |
| | F3-F8 | 88.28*** | 0.96 | 8.81e-05 |
| | Fz-F4 | -97.55*** | -1.60 | .000297 |
| | Fz-Fp2 | 60.84** | 0.83 | .001631 |
| | F4-Fp1 | 92.22* | 1.36 | .010941 |
| | F4-F7 | -50.82* | -0.57 | .019071 |
| | Fp1-T4 | -22.93 | -0.24 | .190731 |
| ANAM-4 | (Intercept) | 0.90 | | .111268 |
| | C3-F8 | -12.64*** | 0.46 | .000127 |
| | F4-T3 | 9.02** | -0.49 | .001723 |

| | | | |
|---|---|---|---|
| C4-T4 | 4.94** | -0.15 | .006325 |
| Fp2-F7 | -4.38** | 0.14 | .004966 |

*b* represents the unstandardized regression weights, and *beta* indicates the standardized regression weights. *p* is the p-value of the predictor in the regression model presented in the table. * $p < .05$,** $p > .01$, *** $p > .001$.
